# Identification of previously untypable RD cell line isolates and detection of EV-A71 genotype C1 in a child with AFP in Nigeria

**DOI:** 10.1101/334094

**Authors:** M.O. Adewumi, T.O.C. Faleye, C.O. Okeowo, A.M. Oladapo, J. Oyathelemhi, O.A. Olaniyi, O.C. Isola, J.A. Adeniji

## Abstract

We previously attempted to identify 96 nonpolio enteroviruses (EVs) recovered on RD cell culture from children <15 years with acute flaccid paralysis (AFP) in Nigeria. We succeeded in identifying 69 of the isolates. Here, we describe an attempt to identify the remaining 27 isolates.

Twenty-six (the 27^th^ isolate was exhausted) isolates that could not be typed previously were further analyzed. All were subjected to RNA extraction, cDNA synthesis, enterovirus 5‟-UTR– VP2 PCR assay and a modified VP1 snPCR assay. Both the 5’-UTR – VP2 and VP1 amplicons were sequenced, isolates identified and subjected to phylogenetic analysis.

Twenty of the 26 isolates analyzed were successfully identified. Altogether, 23 EV strains were recovered. Thesebelong to 11 EV (one EVA, nine EVB and one EVC) types which were EVA71 genotype C1 (1 strain), CVB3 (7 strains), CVB5 (1 strain), E5 (2 strain), E11 (3 strains), E13 (2 strain), E19 (1 strain), E20 (1 strain), E24 (2 strains), EVB75 (1 strain) and EVC99 (2 strains). Of the 11 EV types, the 5’-UTR-VP2 assay identified seven while the VP1 assay identified 10. Both assays simultaneously detected 7 of the 11 EV types identified in this study with 100% congruence.

In this study we identified 20 of 26 samples that were previously untypable. In addition, we provided evidence that suggests that a clade of EVA71 genotype C1 might have been circulating in sub-Saharan Africa since 2008. Finally, we showed that the 5’-UTR-VP2 assay might be as valuable as the VP1 assay in EV identification.

## INTRODUCTION

Enteroviruses are members of the genus Enterovirus in the Picornaviridae family. Within the genus are 13 species and over 100 serotypes distributed into the different species [1]. The best studied members of the genus are the polioviruses (PVs) which are serotypes (1, 2 and 3) in Species C. Enteroviruses have an ∼7.5kb, positive-sense, single stranded RNA genome with a single open reading frame (ORF). The ORF is flanked on both sides by untranslated regions (UTRs). The 5’-UTR has subregions that are conserved among all enterovirus types hence it‟s being used in Panenterovirus detection assays [2, 3]. The 3‟-UTR, on the other hand, has sub-regions that are conserved within species and is consequently used for EV species determination [4]. Determination of EV serological types were doneusing neutralization assays[2, 3, 5]. However, since a correlation was established between serological types of enteroviruses and the sequences of the VP1 gene [5], it has been used for enterovirus identification.

In most developing nations globally, Poliovirus surveillance is the only enterovirus surveillance system in existence. Surveillance for PV entails looking for the virus in sewage and in children presenting with Acute Flaccid Paralysis (AFP) [6]. In line with the WHO recommendation, the global poliovirus surveillance programme uses RD (from Human Rhabdomyosarcoma [7]) and L20b (Mouse L cell line genetically modified to express the human poliovirus receptor [8, 9]) cell lines for poliovirus isolation [3, 4, 6]. While L20b cell line seems to be more specific for the polioviruses, many non-polio enteroviruses (NPEVs) also replicate in RD cell line [3, 4, 6]. Consequently, the global PV surveillance programme, generates as a by-product several NPEVs which till date provide most of the information that exist on NPEV diversity.

In Nigeria, there have been two studies [10, 11] that have investigated EV genetic diversity in AFP cases. While the first study [10] investigated samples dating from 2002 to 2004, the second [11] investigated samples collected in 2014 (about a decade after the first study). In our 2014 study [11], there were 27 isolates recovered on RD cell line from AFP cases that we could not determine their serotypes. This study was consequently designed to use alternative strategies to determine the serotypes of these previously untypables isolates.

## METHODOLOGY

### Samples

The 27 samples that remained untypable from our previous study [11] were the subject of this study. However, because, one of the samples was already exhausted, it could not be further analyzed. Hence, only the remaining 26 samples were further analyzed. To be precise, these 26 isolates were recovered on RD cell line from children (<15 years old) presenting with AFP in Nigeria in 2014 [11]. The samples had been previously subjected to three enterovirus screens (Figure 1 and [11]) but, their respective types remained to be determined.

**Figure 1:**
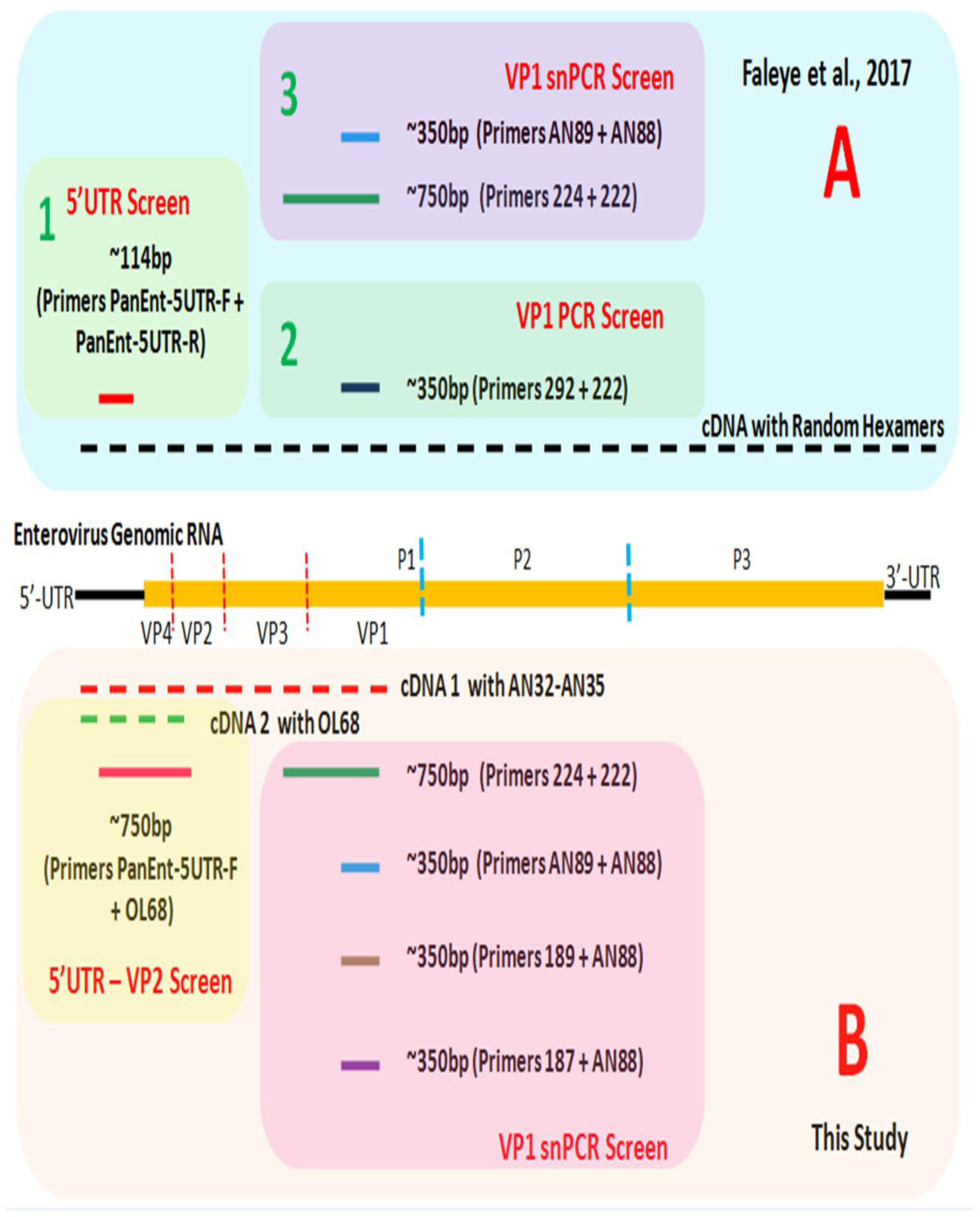
The algorithm followed in this study. Section "A‟ of the figure depicts the screens the isolates were previously subjected to in Faleye et al., 2017 [11]. Section "B‟ shows the alternative approach deployed for identifying the isolates in this study.

### RNA extraction and cDNA synthesis

All isolates were subjected to RNA extraction using the Total RNA extraction kit (Jena Bioscience, Jena, Germany) as instructed by the manufacturer. Subsequently, cDNA was made using the SCRIPT cDNA synthesis kit (Jena Bioscience, Jena, Germany) following manufacturer‟s instructions with slight modifications. Two (2) separate cDNA sets were made (Figure 1). Rather than use Random Hexamers as described in [11], Primers AN32 – AN35 were used for synthesis of cDNA 1 as previously described [6, 12, 13] while primer OL68 was used for synthesis of cDNA 2 [14].

### Polymerase Chain Reaction (PCR) assays

Two different PCR assays were done in this study. Assay 1 amplified a ∼750bp amplicon spanning part of the 5’-UTR through VP4 to the 5‟-end of VP2 (5’-UTR – VP2). Primers PanEnt-5UTR-F and OL68 [14] alongside cDNA two were used for this assay.

The second PCR assay (assay 2) is the modified version of the Nix et al., [12] semi-nested PCR assay described in Adeniji et al., [15]. The first round PCR amplifies a ∼750bp amplicon spanning parts of VP3 and VP1 (VP3 – VP1). Primers 224 and 222 [12] alongside cDNA1 were used for this assay. Three second round PCR assays, each amplifying a ∼350bp amplicon within the 5‟ part of VP1 were done as described in Adeniji et al., [15]. The three second round assays (PE, EV-A/C and EV-B) used forward primers AN89, 189 and 187, respectively, paired with same reverse primer AN88. The product of the first round PCR assay was used as template for all the three second round assays.

Thermal cycling was done using a Veriti Thermal cycler (Applied Biosystems, California, USA). For assay one and first round of assay two, cycling conditions were 94°C for 3 minutes, then 45 cycles of 94°C for 30 seconds, 42°C for 30 seconds, and 60°C for 60 seconds, with ramp of 40% from 42°C to 60°C. This was then followed by 72°C for 7 minutes and held at 4°C until the reaction was terminated. Cycling conditions for second round of assay two were similar to that of the first round except for the extension time that was reduced to 30 seconds. PCR products of assay 1 and the second round PCR product of assay 2 were resolved on a 2% Agarose gel stained with ethidium bromide and viewed by ultraviolet (UV) light on a transilluminator.

### Sequencing and Identification

All amplicons of PCR reactions with the expected band size were shipped to Macrogen Inc, Seoul, South Korea, where amplicon purification and sequencing were done. Sequencing was performed using the forward and reverse primers for each of the respective assays. Subsequently, the sequence data was used for enterovirus genotype and species determination. For assay 1 the sequence data was subjected to the NCBI BLASTn tool while the sequence data for the second round PCR product of assay 2 were subjected to the enterovirus genotyping tool [16].

### Phylogenetic Analysis

Using the default settings of the CLUSTAL W program in MEGA 5 software [17] sequences of the EVA71 described in this study were aligned alongside those retrieved from GenBank. Afterwards a neighbor-joining tree was constructed using the same MEGA 5 software with the Kimura-2 parameter model [18] and 1,000 bootstrap replicates.

### Nucleotide sequence accession numbers

All the sequences reported in this study have been deposited in GenBank and assigned accession numbers MH379115-MH379135 and MH397260-MH397268.

## RESULTS

### PCR assays

Of the 26 samples subjected to the 5’-UTR-VP2 screen, amplicons were recovered from 11. The remaining 15 samples were negative for the screen. For the three VP1 snPCR screens, 19, 4 and 18 of the 26 samples subjected to these assays were positive for the PE, EV-A/C and EV-B assays, respectively. (Table 1).

**Table 1:** Results of the different PCR assays and identities of isolates typed.

| S/N | 5'-UTR –<br>VP2<br>ASSAY |  | VP1 ASSAYS |  |  |  |  |  | Serotypes | Number<br>of<br>Isolates |
| --- | --- | --- | --- | --- | --- | --- | --- | --- | --- | --- |
|  |  |  | AN89 (PE) |  | 189 (EV-A/C) |  | 187 (EV-B) |  |  |  |
|  | PCR | SEQ | PCR | SEQ | PCR | SEQ | PCR | SEQ |  |  |
| 1 | + | E24 | + | E24 |  |  | + | E24 | E24 | 1 |
| 2 |  |  |  |  |  |  |  |  |  |  |
| 3 | + | E24 | + | E24 |  |  | + | E24 | E24 | 1 |
| 4 |  |  | + | CVB3 |  |  |  |  | CVB3 | 1 |
| 5 | + | E19 | + | E19 |  |  | + | E19 | E19 | 1 |
| 6 | + | E5 | + | E5 |  |  | + | E5 | E5 | 1 |
| 7 |  |  | + | E5 |  |  | + | E5 | E5 | 1 |
| 8 |  |  | + | CVB3 |  |  | +W | UnD | CVB3 | 1 |
| 9\$ | + | CVB3 | | | | | | | CVB3 | 1 |
| 10 | + | E20 | + | E20 | + | E20 | + | E20 | E20 | 1 |
| 11 | + | CVB5 | + | CVB5 |  |  | +W | CVB5 | CVB5 | 1 |
| 12 |  |  | + | CVB3 |  |  | +W | CVB3 | CVB3 | 1 |
| 13 |  |  | + | CVB3 |  |  | + | CVB3 | CVB3 | 1 |
| 14 |  |  |  |  |  |  | +W | CVB3 | CVB3 | 1 |
| 15 | + | CVB3 | + | CVB3 |  |  | + | CVB3 | CVB3 | 1 |
| 16 | +W | UnD |  |  |  |  |  |  |  |  |
| 17 |  |  | + | E11 |  |  | +W | UnD | E11 | 1 |
| 18\$ | + | EVB75 | | | | | +W | UnD | EVB75 | 1 |
| 19 |  |  | + | NS* | + | UnD |  |  |  |  |
| 20 |  |  | + | UnD |  |  | +W | UnD |  |  |
| 21 |  |  | + | EVC99 | + | EVC99 |  |  | EVC99 | 1 |
| 22 |  |  |  |  |  |  |  |  |  |  |
| 23 |  |  | + | EVC99 |  |  | + | E11 | EVC99/E11 | 2 |
| 24 |  |  |  |  |  |  |  |  |  |  |
| 25 |  |  | + | E11 |  |  | + | E13 | E11/E13 | 2 |
| 26 | +W | UnD | + | E13 | + | EVA71 | + | NS* | E13/EVA71 | 2 |
|  | 11 | 9 | 19 | 17 | 4 | 3 | 18 | 13 |  | 23 |

W = Weak; UnD = Unusable Data; E = Echovirus; EV = Enterovirus; CV = Coxsackievirus; NS\* = Not Sequenced; \$ = Identified by only the 5'-UTR – VP2 assay

### Sequencing and Identification

All the 11amplicons recovered from the 5’-UTR-VP2 screen were sequenced. However, only nine (9) of the sequence data generated were useable. The remaining two (2) were not useable due to the presence of multiple peaks. A BLASTn search of GenBank typed the EVs as Echovirus (E) 5 (1 strain), E19 (1 strain), E20 (1 strain), E24 (2 strains), Coxsackievirus (CV) B3 (2 strains), CVB5 (1 strain) and EVB75 (1 strain) (Table 2).

**Table 2:** Identification of Enterovirus types using the 5’-UTR-VP2 sequence data and a BLASTn search of GenBank

| S/N | SAMPLE ID | ACCESSION NUMBER OF MOST SIMILAR SEQUENCE | IDENTITY | E VALUE | QUERY COVER | ENTEROVIRUS TYPE |
| --- | --- | --- | --- | --- | --- | --- |
| 1 | 1 | KP036484.1 | 86% | 0.0 | 99% | E24 |
| 2 | 3 | KP036484.1 | 86% | 0.0 | 99% | E24 |
| 3 | 5 | KY792585.1 | 85% | 0.0 | 100% | E19 |
| 4 | 6 | HM775882.1 | 82% | 9e-174 | 100% | E5 |
| 5 | 9 | KC481610.1 | 83% | 1e-172 | 99% | CVB3 |
| 6 | 10 | KF812551.1 | 87% | 0.0 | 100% | E20 |
| 7 | 11 | KT285014.1 | 88% | 0.0 | 100% | CVB5 |
| 8 | 15 | KR107057.1 | 85% | 0.0 | 100% | CVB3 |
| 9 | 18 | AY556070.1 | 86% | 0.0 | 100% | EVB75 |

Only 18 of the 19 amplicons recovered from the PE screen were sequenced. The 19^th^ amplicon was not sequenced because, it leaked during transit (from Nigeria to South Korea) to the sequencing facility. Of the 18 amplicons sequenced, 17 were useable. The remaining one (1) was not useable due to the presence of multiple peaks. The Enterovirus Genotyping Tool (EGT) identified the 17 as E5 (2 strains), E11 (2 strains), E13 (1 strain), E19 (1 strain), E20 (1 strain), E24 (2 strains), Coxsackievirus (CV) B3 (5 strains), CVB5 (1 strain) and EVC99 (2 strains) (Table 1).

All the four (4) amplicons recovered from the EV-A/C screen were sequenced. However, three (3) of the sequence data generated were useable. The remaining one (1) was not useable due to the presence of multiple peaks. The EGT identified the three as E20 (1 strain), EVC99 (1 strain) and EVA71 genotype C1 (1 strain) (Table 1).

Only 17 of the 18 amplicons recovered from the EV-B screen were sequenced. The 18^th^ amplicon was not sequenced because, it leaked during transit (from Nigeria to South Korea) to the sequencing facility. Of the 17 amplicons sequenced, 13 were useable. The remaining four (4) were not useable due to the presence of multiple peaks. The Enterovirus Genotyping Tool (EGT) identified the 13 as E5 (2 strains), E11 (1strain), E13 (1 strain), E19 (1 strain), E20 (1 strain), E24 (2 strains), Coxsackievirus (CV) B3 (4 strains) and CVB5 (1 strain) (Table 1).

In all, 20 of the 26 isolates where identified as belonging to 11 (one EV-A, nine EV-Bs and one EV-C) EV types. Also, 23 different enterovirus strains were identified in the 20 isolates typed. Precisely, the 23 different enterovirus strains were EVA71 genotype C1 (1 strain), CVB3 (7 strains), CVB5 (1 strain), E5 (2 strains), E11 (3 strains), E13 (2 strains), E19 (1strain), E20 (1 strain), E24 (2 strains), EVB75 (1 strain) and EVC99 (2 strains) (Table 3).

**Table 3:**
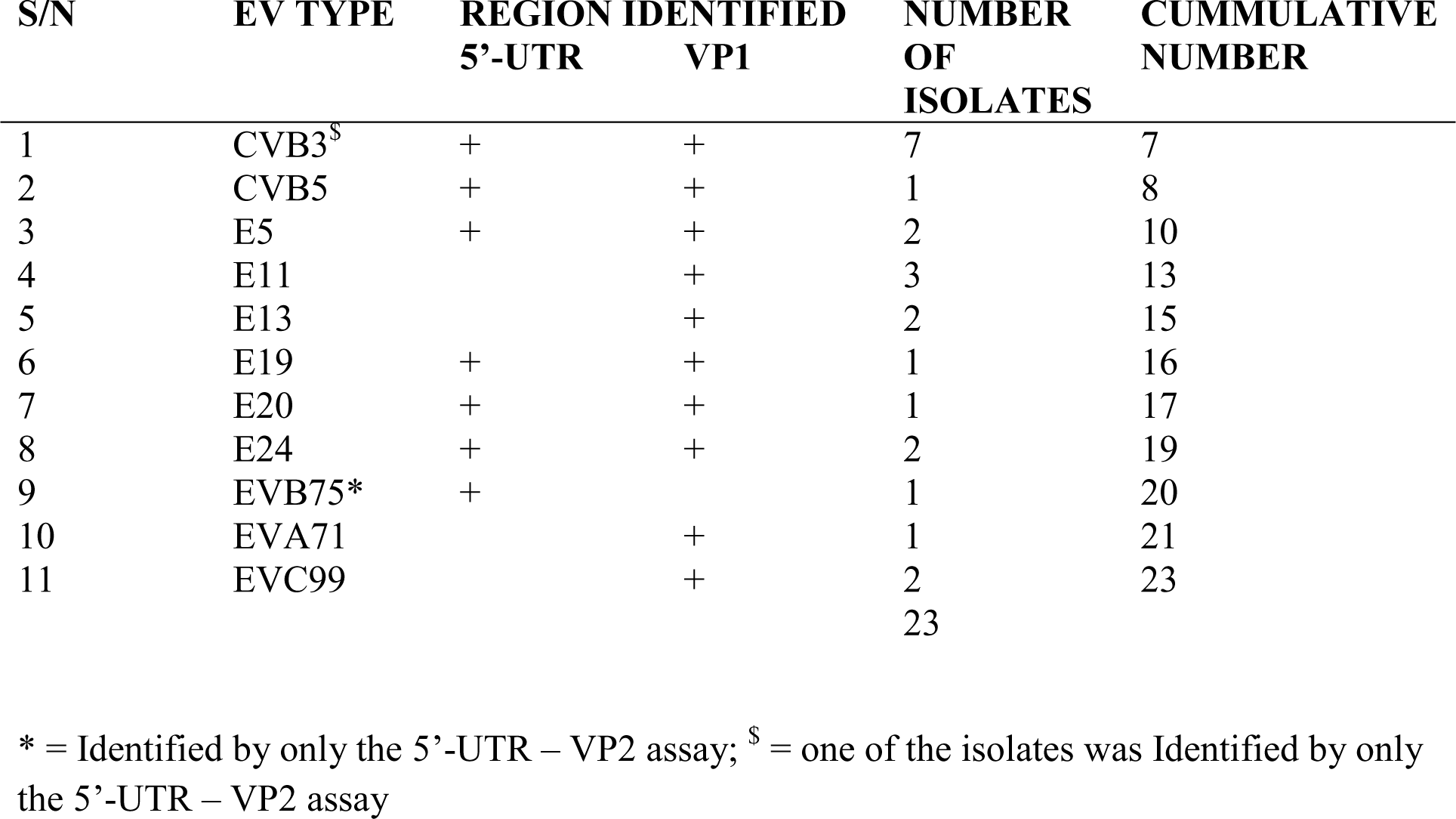
Enterovirus types identified in this study, the genomic region used for identification and number of isolates of each type.

### Comparison of the 5’-UTR-VP2 and the VP1 screens

Of the 11 EV types identified in this study, the 5’-UTR-VP2 assay was able to identify seven (7) of them while the VP1 assay identified 10 of them (Table 3). Both assays simultaneously detected seven (7) of the 11 EV types identified in this study (Table 4). EVB75 was only identified by the 5’-UTR-VP2 assay while E11, E13, EVA71 and EVC99 were only identified by the VP1 assay (Table 3).

**Table 4:** Isolates typed using both 5’-UTR-VP2 and VP1 genomic regions

| **S/N** | **ISOLATE ID** | **IDENTITY BY 5'-UTR-**<br>**VP2** | **IDENTITY BY VP1** | **COMMENTS** |
| --- | --- | --- | --- | --- |
| 1 | 1 | E24 | E24 | CONGRUENT |
| 2 | 3 | E24 | E24 | CONGRUENT |
| 3 | 5 | E19 | E19 | CONGRUENT |
| 4 | 6 | E5 | E5 | CONGRUENT |
| 5 | 10 | E20 | E20 | CONGRUENT |
| 6 | 11 | CVB5 | CVB5 | CONGRUENT |
| 7 | 15 | CVB3 | CVB3 | CONGRUENT |

### Phylogenetic Analysis

Only the EVA71 described in this study was subjected to phylogenetic analysis because of the association of EVA71 (and especially Genotype C strains) with neurological manifestations [19]. The tree (Figure 2) confirms that the EVA71 described in this study belongs to genotype C1. It further confirms the result of a BLASTn search (data not shown) that the EVA71 described in this study is most similar to that detected in a child with AFP in Cameroon in 2008 [20].

**Figure 2:**
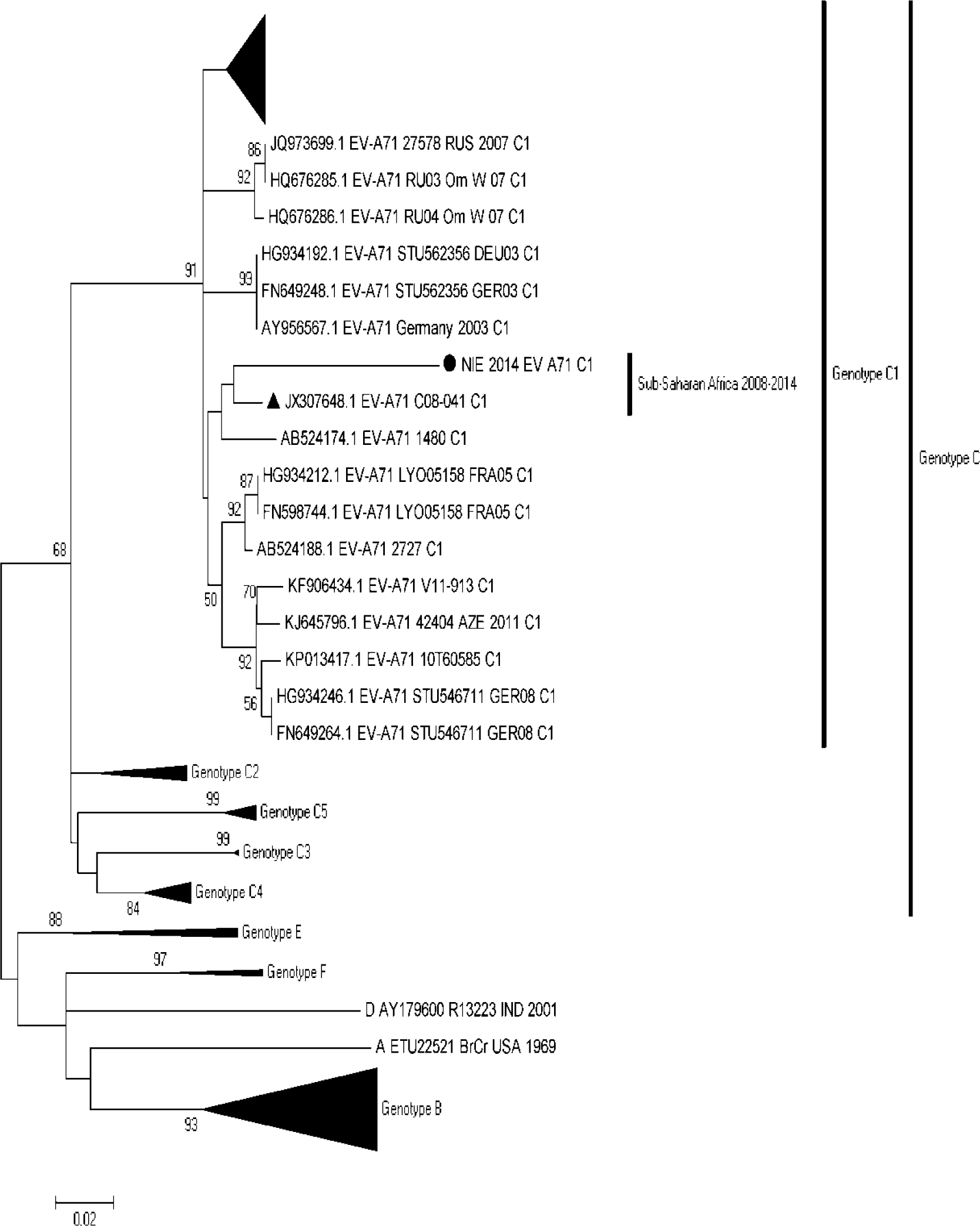
Phylogram of genetic relationship between VP1 nucleotide sequences of EV-A71. The phylogenetic tree is based on an alignment of the partial VP1 sequences. The reference EVA71 sequences described in Bessaud et al., [21] was used as baseline. Added to it was the EVA71 described in this study alongside the top 50 hits of a BLASTn search using the EVA71 described in this study as query. The newly sequenced strain is indicated with black circle. The only other genotype C1 strain detected and described in Africa till date is indicated with a black triangle. Bootstrap values are indicated if >50%.

## DISCUSSION

In our 2014 study [11], there were 27 isolates recovered on RD cell line from AFP cases that we could not determine their serotypes. This study was aimed at using alternative strategies to determine the serotypes of these previously untypables isolates. At the commencement of this study we had access to 26 of the samples (the 27^th^ was already exhausted). We successfully identified 20 of the 26 samples analyzed. From the 20 isolates, 23 EV strains were recovered, 20 of which were EV-Bs (Table 1). This is not surprising in the light of the EV-B bias of RD cell line [20, 22].

Of the 23 EV strains typed in this study, 21 were unambiguously identified using the VP1 assay (Table 1). As shown in Figure 1, two basic differences exist between the VP1 assay used in this study and that used in Faleye et al., [11]. The first difference lies in the primers used for cDNA synthesis. As opposed to the previous study [11] where cDNA was done using Random Hexamers, cDNA for the VP1 assay was made using the primers AN32-AN35 [6, 12]. This obviously, significantly improved our capacity to identify some of the previously untypable EVs. In fact, 17 of the 21 strains detected by the VP1 assay can be accounted for by this modification in the assay design. This therefore confirms, the sensitivity of this cDNA synthesis algorithmand its consequent superiority to using random Hexamers for EV VP1 amplification.

The second difference between the VP1 assay used in this study and that used in Faleye et al., [11], lies in the second round PCR assays. While only one second round PCR assay was done in Faleye et al., [11], three different second round PCR assays were done in this study (Figure 1). We had previously shown that this modification increases the sensitivity of the assay and especially its capacity for EV co-infection resolution [15, 16, 24]. In this study, four (4) of the 21 strains detected by the VP1 assay can be accounted for by this modification in the assay design. Of the four strains detected exclusively by this modification, the CVB3 in Sample 14 (Table 1) would have been completely missed if not for this assay modification. This shows that a sample declared negative by the AN89-AN88 (PE) assay might not be truly negative. Considering the very variable nature of the VP1 protein, it is possible that the virus in the sample of interest might have mutations in the AN89 primer binding site that makes it impossible for the primer to bind to the site.

The other three strains detected exclusively by this modification (three different second round PCR assays; Figure 1) were cases of co-infection (Table 1). To be precise, the E11, E13 and EVA71-C1 detected in samples 23, 25 and 26 would have also been missed should we have relied on the PE assay alone for EV VP1 amplification. Considering how common EV co-infection is (3/26; in this study), coupled with their role in cVDPV emergence [25, 26], we have repeatedly [13, 15, 16, 24] emphasized the need to employ co-infection resolution assays when prospecting for enteroviruses. Against this backdrop, it is important to bear in mind any EV surveillance study that uses the PE (AN89 + AN88) assay alone for EV VP1 detection and subsequently, EV identification might not be capturing the complete picture of EV diversity in the samples screened.

This modified protocol enabled us to detect the EVA71 genotype C1 for the first time in Nigeria. Prior this study, only genotype E of EVA71 had been described in the country [10, 27]. Both a BLASTn search (data not shown) of the GenBank and phylogenetic analysis (Figure 2) suggests that the EVA71 genotype C1 described in this study is most similar to one recovered from a child with AFP in Cameroon in 2008 [20]. Though EVA71 genotype Cs have been associated with clinical manifestations that might have neurological complication, and both EVA71 genotype C1 strains described in sub-Saharan Africa till date were from children with AFP, it might be premature to conclude they were responsible for the associated clinical presentation. For example, firstly, in the Nigeria case described in this study, if not for the co-infection resolution modification to the VP1 assay, only E13 would have been detected in that isolate. In the light of co-infection with both E13 and EVA71 genotype C1 in this case, to which of the two EV types do we ascribe the clinical presentation? Of course, the synergistic effect of EV co-infections cannot be overlooked, but until rigorous experimental data is provided to document such associations, it is only reasonable that all be interpreted with caution. Especially those cases in which no attempt was made to detect or resolve possible co-infections.

Furthermore, the topology of the EVA71 tree suggest that the clade detected in Cameroon in 2008 and subsequently in Nigeria in 2014 might have been circulating between that time (and if the regional confinement hypothesis is anything to go by) in the region. Whether this clade is still circulating is not known. However, this finding further necessitates the need to rigorously search for the virus in the population.

With respect to the 5’-UTR-VP2 assay, of the 23 EV strains typed in this study, 9 were identified using this assay (Table 2). It is however important to note that seven (7) of the nine EV types identified by the 5’-UTR-VP2 assay were also simultaneously identified by the VP1 assays and their identities by both assays were congruent (Table 4). The remaining two isolates only identified using the results of the 5’-UTR-VP2 assay was because the VP1 assays failed to work on these isolates. Hence, there was no VP1 data to compare (Tables 1 and 3). However, considering the 100% congruence of those for which both 5’-UTR-VP2 and VP1 data exist (Table 4), we feel comfortable taking the identification as correct as several other studies have shown [14, 28]. Our findings therefore also demonstrate the usefulness of this assay for EV identification.

## CONCLUSIONS

In this study we identified 20 of 26 samples that were previously untypable [11]. We showed that modifying the algorithm used in Faleye et al., 2017 to include AN32-AN35 specific cDNA synthesis and expanding the second round PCR assay to accommodate co-infection resolution can significantly increase the sensitivity of the EV VP1 assay. We further provided evidence that suggest that a clade of EVA71 genotype C1 might have been circulating in sub-Saharan Africa since 2008. Finally, we showed that the 5’-UTR-VP2 assay might be as valuable as the VP1 assay in EV identification.

## ACKNOWLEDGEMENTS

We thank the WHO National Polio Laboratory in Ibadan, Nigeria for providing the anonymous isolates analyzed in this study. This study was funded by contributions from authors.

## AUTHOR CONTRIBUTIONS

1. Study Design (AMO,FTOC, AJA)

2. Sample Collection, laboratory and data analysis (All Authors)

3. Wrote the first draft of the Manuscript (AMO, and FTOC)

5. Revised the Manuscript (All Authors)

6. Read and Approved the Final Draft (All Authors)

7. AJA supervised this study

